# The functional role of episodic memory in spatial learning

**DOI:** 10.1101/2021.11.24.469830

**Authors:** Xiangshuai Zeng, Laurenz Wiskott, Sen Cheng

**Affiliations:** Institute for Neural Computation, Faculty of Computer Science, Ruhr University Bochum

## Abstract

Episodic memory has been studied extensively in the past few decades, but so far little is understood about how it drives behavior. Here we propose that episodic memory can facilitate learning in two fundamentally different modes: retrieval and replay. We study their properties by comparing three learning paradigms using computational modeling based on visually-driven reinforcement learning. Firstly, episodic memory is retrieved to learn from single experiences (one-shot learning); secondly, episodic memory is replayed to facilitate learning of statistical regularities (replay learning); and, thirdly, learning occurs online as experiences arise with no access to past experiences (online learning). We found that episodic memory benefits spatial learning in a broad range of conditions, but the performance difference is meaningful only when the task is sufficiently complex and the number of learning trials is limited. Furthermore, the two modes of accessing episodic memory affect spatial learning distinctly. One-shot learning is initially faster than replay learning, but the latter reaches a better asymptotic performance. Our model accounts for experimental results where replay is inhibited, but the hippocampus, and hence episodic memory, is intact during learning. Understanding how episodic memory drives behavior will be an important step towards elucidating the nature of episodic memory.

## Introduction

Even though there is widespread consensus that episodic memory (EM) is the memory of personally experienced episodes (Tulving, 1972), the precise conceptualization of episodic memory has been difficult to come by. One source of this difficulty might be a dominant focus on the properties of EM, whereas little is known about its function. It seems intuitive that information from past experiences is useful somehow, but how exactly does the stored experience of a particular past episode drive learning and future behavior?

Research on human memory often addresses this question from an abstract, conceptual perspective instead of a mechanistic and computational one. For instance, Klein et al. (2009) suggested that maintaining a pool of episodic memories enables its owner to reevaluate an individual’s past behavior in light of new information, thus serving an important role in social interaction. Mahr and Csibra (2017) analyzed the communicative function of episodic memory from a philosophical point of view and argued that episodic memory plays a generative role in the justification of our beliefs about past events. Another influential idea is based on the survival processing benefit, which refers to the fact that subjects can remember names of objects that are relevant for survival in the wilderness better than non-relevant words (Nairne et al., 2007). The adaptive memory theory argues that episodic memory has adapted to ensure our survival in the kind of environments that our stone-age ancestors lived in (Nairne and Pandeirada, 2016). However, EM probably has other functions beyond remembering survival-relevant items that are not covered by the adaptive memory theory. Suddendorf and Corballis (1997, 2007) suggested more broadly that mentally traveling into the past is an epiphenomenon of the capacity to mentally travel into the future. Forecasting the future, they argued, enables us to take the appropriate actions in the present to ensure a favorable outcome in the future. While each of the above hypotheses suggests a potential function of EM, none spells out how the recalled memory drives upcoming or future behavior.

In computational neuroscience, a specific suggestion is that EM provides the data to extract regularities from multiple, repeated experiences (Nadel and Moscovitch, 1998; Cheng, 2017). The Complementary Learning Systems (CLS) theory makes a related suggestion and postulates that replay of episodic memory supports the integration of novel information into an existing semantic network (McClelland et al., 1995; Kumaran et al., 2016). According to CLS theory, replay facilitates interleaved training, i.e., the alternating presentation of novel and old information, to avoid catastrophic interference (McCloskey and Cohen, 1989). Although hippocampal replay was hypothesized to play a role in learning more than three decades ago (Buzsaki, 1989), and CLS theory provides a specific suggestion for its computational function, it is still lacking a functional role of EM in driving behavior, which is required for a measure of performance. Such a link is provided by reinforcement learning (RL) studies in which agents need to take sequences of actions in an environment to maximize the expected accumulated reward. Early work used online learning exclusively, i.e., an experience drove learning exactly once. Later it was found that replaying earlier experiences speeds up learning in many RL tasks (Lin, 1992). Recent advances in utilizing episodic-like memory led to human-level performance on many (video) games (Mnih et al., 2015). However, even though replay in these technical applications improved performance, it has not been studied what this implies about the functional role of EM in biological settings.

In this paper, we use algorithms developed in the framework of RL to quantitatively study and compare two different operating modes in which the mammalian brain could use EM in spatial learning. we contrast two paradigms which use EM in different ways, i.e. retrieval and replay, and one paradigm which does not access EM. We hypothesize that the learning curves of the three paradigms will show characteristic differences. We focus on spatial learning because it is strongly linked to the hippocampus (Broadbent et al., 2004), much is known about hippocampal activity during spatial navigation and learning (Moser et al., 2008), and the hippocampus is closely linked to EM (Tulving and Markowitsch, 1998). Hence, spatial learning offers a wide range of experimental results to compare with. Our simulations were divided into two parts. First, we simulated the three learning paradigms separately to test our hypothesis and analyzed their individual characteristics. The three learning paradigms solve spatial learning tasks at different speeds, and the harder the task is, the more profound the difference is. The agents also show different patterns of behaviour and reach different asymptotic performance during and at the end of the learning. Second, we conducted simulations where the three paradigms coexist and interact with each other in an agent, which might be a more realistic model of how animals learn in spatial navigation. Consistent with experimental findings (Girardeau et al., 2009; Ego-Stengel and Wilson, 2010), disrupting hippocampal replay slows down learning in our model. Our results lead to predictions about the nature and functions of episodic memory in spatial learning.

### Hypotheses

One key to understanding the function of EM is to recognize that EM can provide information that is useful for learning in at least two fundamentally different modes. Firstly, the most direct way of using EM to drive behavior is to retrieve a sequence of events that composes the episode and that includes information about the actions performed, and use that information directly to learn a sequence of actions. For instance, a rat might go to one arm of a T-maze because it remembers that it found a piece of cheese in that arm once before (Fig. 1A). We term this kind of learning *one-shot learning*. Secondly, EM can be replayed offline repeatedly to drive learning in the neocortex, as suggested before (McClelland et al., 1995; Nadel and Moscovitch, 1998; Cheng, 2017). Replay enables the neocortex to acquire the information about how to solve a cognitive task from multiple memory traces of similar experiences. For instance, a rat running in a T-maze might learn that a piece of cheese is always located in the right arm. We term this kind of learning *replay learning*. Furthermore, learning can also occur without employing EM, since the neocortex can directly extract semantic information from online experiences as they occur (*online learning*). By semantic information we mean statistical information about objects and their relationships, which includes symbolic or linguistic representations, but might be sub-symbolic as well (Cheng et al., 2016; Cheng, 2017). Unlike in *replay* learning, experiences are not stored during online learning and can be used only once to effect changes in the neural network that drives behavior.

**Figure 1:**
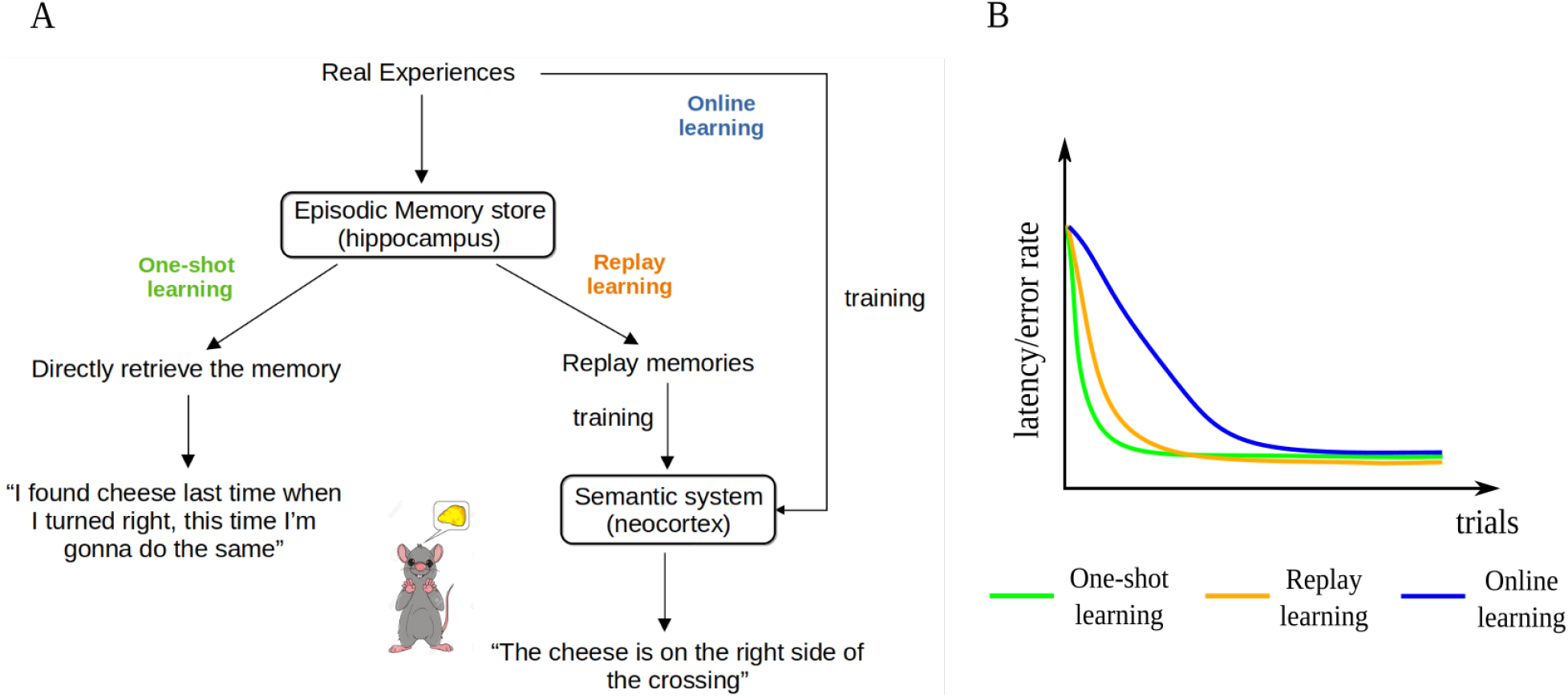
The three hypothesized learning paradigms and their learning curves. **A**: A schematic illustration of the three learning paradigms for the example of a rat running in a T-maze. **B**: The hypothesized learning curves of the three learning paradigms for a spatial learning task.

We hypothesize that the learning speeds of the three learning paradigms have the relationship: one-shot learning *>* replay learning *>* online learning. The direct use of episodic memory does not require multiple updates so it is one-shot and depends on the hippocampus (Cheng, 2013). By contrast, it takes time for the neocortex, an extended and distributed network, to extract semantic information from memory replay (Cheng, 2017), so we hypothesize that replay learning is slower than one-shot learning. However, it is still faster than online learning, because replaying previous experiences multiple times increases the number of exposures and the interleaved training overcomes interference among different memories (McClelland et al., 1995). The three learning paradigms coexist in a healthy brain, whereas hippocampal animals or patients can rely only on online learning. Indeed, experimental studies show that hippocampal patients are still able to acquire new semantic memories but generally require many learning trials to master the new knowledge, while controls learn the same contents after a single trial (O’Kane et al., 2004; Rosenbaum et al., 2005). Similar observations have been made in rodents as well (Wiltgen et al., 2006; Kosaki et al., 2014). These hypotheses are summarized and formalized in the learning curves of the three learning paradigms (Fig. 1B).

In this paper, we investigate the quantitative properties of the three learning paradigms individually in a computational model of spatial learning and describe the interactions of the three paradigms to examine the effect of experimentally abolishing memory replay. These studies allow us to test and confirm our hypotheses about the learning curves of the three learning paradigms.

## Results

To test our hypotheses regarding the function of EM in spatial learning, we employed visually-guided reinforcement learning in a virtual-reality modeling framework (Fig. 2) that was developed to study models of rodent behavior in spatial navigation and extinction learning (CoBeL-RL, Walther et al., 2021). Within this framework, We implemented three reinforcement learning algorithms: Model-free Episodic Control (EC), Deep Q Learning (DQN) and online Deep Q Learning (online DQN) (see Materials and Methods for the details of the algorithms). The agents trained using the three algorithms are referred to as EC agent, DQN agent, and online DQN agent, respectively. The EC agent uses sequences of stored experiences to learn directly from episodic memory, the DQN agent uses experience replay to drive learning in a deep neural network (DNN) and the online DQN agent does not have access to previous experiences (Fig. 3). Therefore, the three agents model the three learning paradigms: one-shot learning, replay learning, and online learning, respectively.

**Figure 2:**
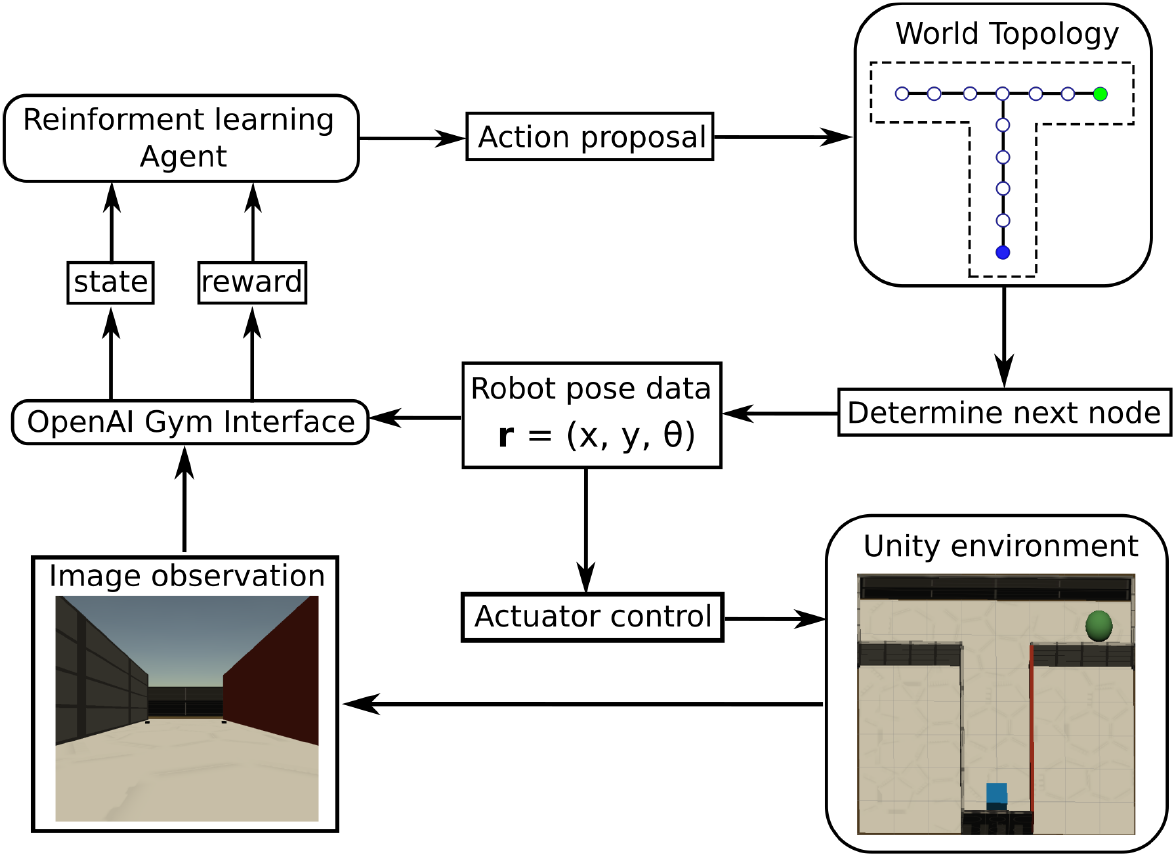
Schematic of the CoBeL-RL simulation framework. Modules are represented by rounded rectangles, data and commands by regular rectangles. The information or control flows are depicted by arrows.

**Figure 3:**
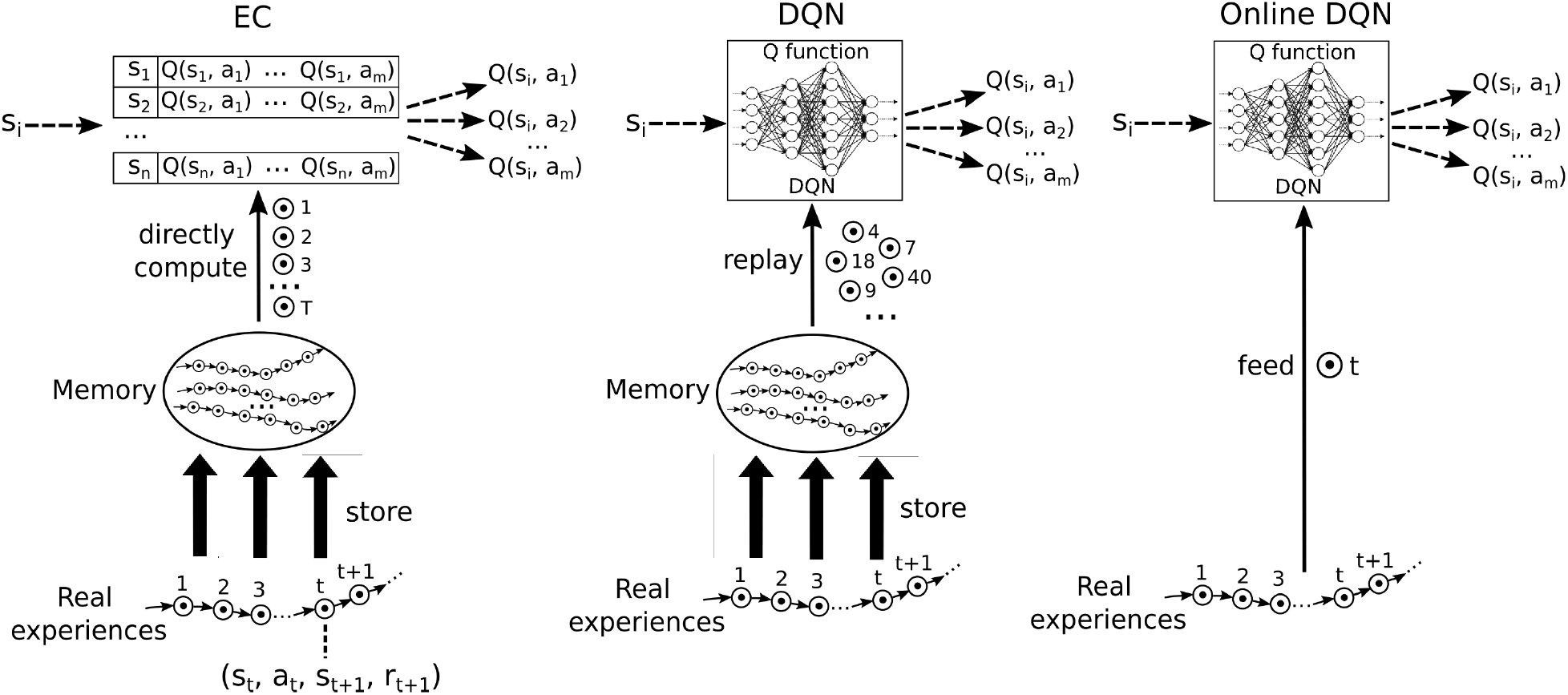
A schematic illustration of the three reinforcement learning algorithms used in this study. Each circle with a dot inside represents an experience, defined as a (state, action, reward, next state) tuples. Real experiences are collected sequentially and stored in memory. Model free episodic control (EC) uses this sequential information to systematically extract reward information, which are stored in a table of Q values. The Deep Q network (DQN) selects experiences randomly for replay to train a deep neural network, which then represents the Q function. The online DQN does not store experiences in memory and uses each experience only once to train the deep neural network representing the Q function.

The three agents had to solve five different spatial navigation tasks (Fig. 4). We first analyzed the learning curves for every environment and then focused on the results from the tunnel maze, because this was where the differences of the three agents became most pronounced. Finally, we conducted simulations of the hybrid agent running in the tunnel maze as a model of how rodents combine the three learning paradigms to learn to navigate and examined the effect of disrupting hippocampal replay.

**Figure 4:**
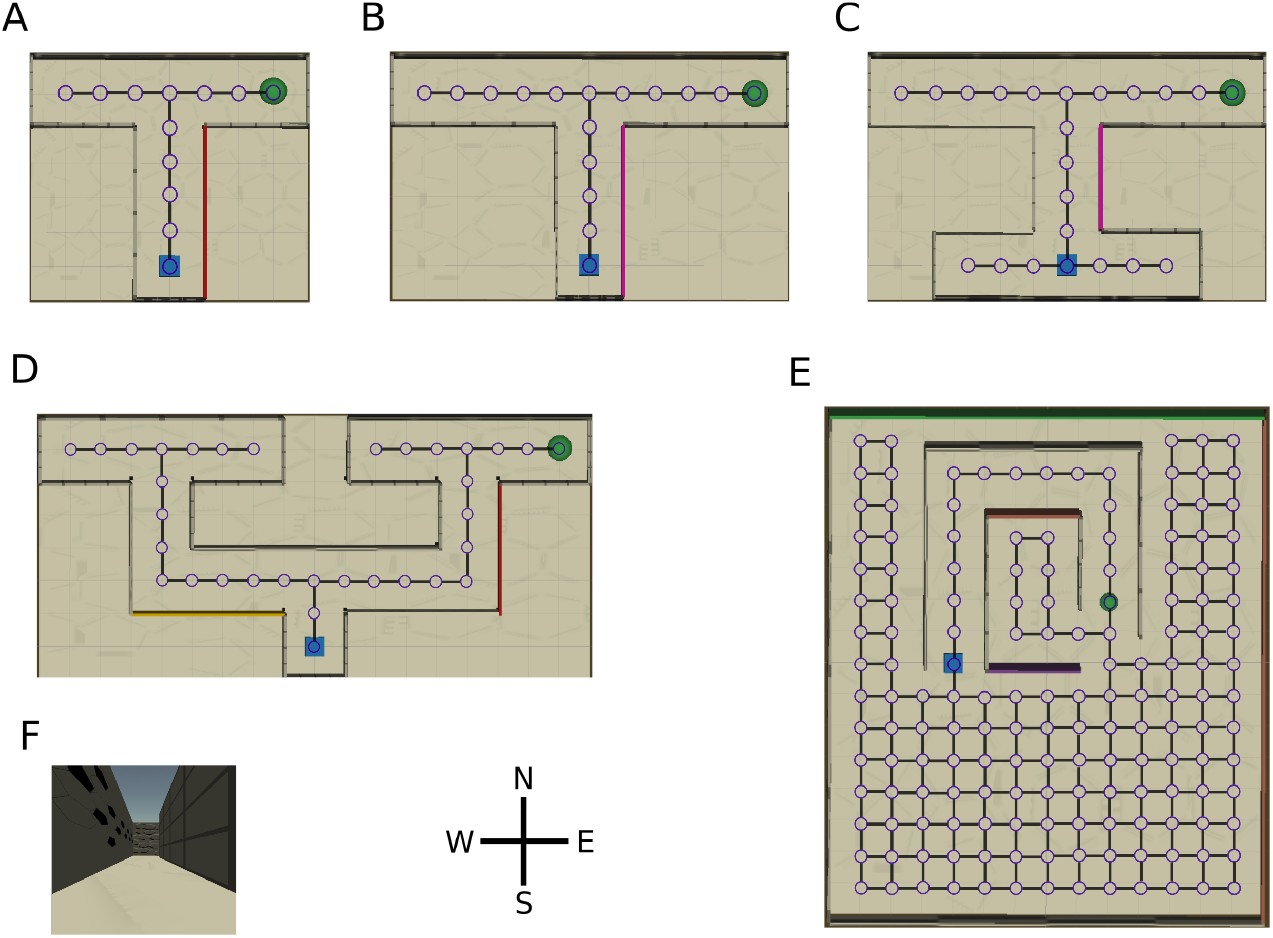
Overview of the virtual environments with their topology graphs. Nodes in the graph represent allowed positions for the agent and solid lines the allowed transitions between positions. At each node, the agent can face in four directions: north, west, south, and east. The starting position of the agent (blue square) and the location of the goal (green disk) remain constant during the simulation. Note, the topology graphs are only drawn on this figure for demonstration purpose, they are not visible to the agent. **A**: T-maze, **B**: long T-maze, **C**: H-maze, **D**: double T-maze, **E**: tunnel maze, and **F**: an example view collected by the camera placed on top of the agent.

### Episodic memory facilitates spatial learning in two different modes

The learning curves for the three agents in the five tasks confirm our hypotheses and reveal interesting details (Fig. 5). Namely, the learning speeds of the three algorithms have the relationship: EC *>* DQN *>* online DQN in all of the tested environments. The learning curves correlate with our intuition about the complexity of the five tasks. For instance, all three agents required a larger initial number of time steps to complete a trial in the double T-maze (Fig. 5D) as compared to the T-maze (Fig. 5A). Also, the learning curves drop more slowly, meaning that more learning trials are required to find the shortest path. These differences in the learning curves are consistent with a higher task complexity of the double T-maze compared to that of the T-maze. By contrast, between the T-maze and the long T-maze, as well as between the double T-maze and the tunnel maze, the learning curves differ only slightly, consistent with the task complexity differing only subtly between the two pairs of mazes.

**Figure 5:**
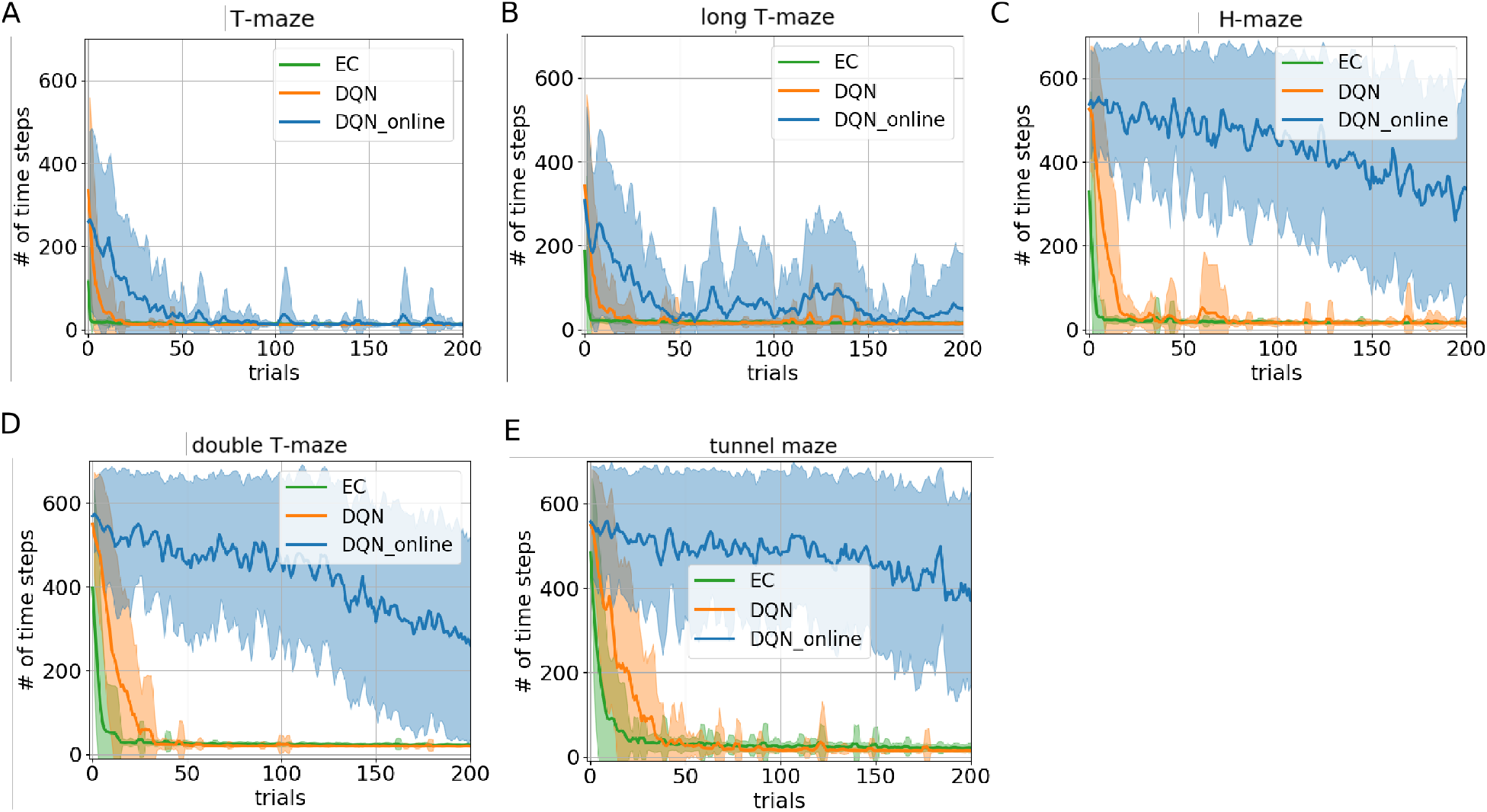
Learning curves for the three learning paradigms in different environments. Each panel shows the number of time steps that the agent takes to find the goal in a specific trial. The curves and shaded areas indicate the mean and the standard deviation, respectively, obtained from 25 independent runs. Only the learning curves for the first 200 trials are shown here because both EC and DQN have converged at this point. Although the curves for online-DQN do not drop much in panel C, D and E, some of the agent still managed to find solutions at the end of the simulation (500 trials). Model-free Episodic Control (EC) represents one-shot learning, Deep Q Learning with memory replay (DQN) represents replay learning, and online Deep Q Learning (online DQN) represents online learning.

Interestingly, the online DQN agent is more sensitive to the change of the task complexity compared to the EC and DQN agents, since its learning curves move up and rightwards dramatically. If we limited the number of trials to 100, then online DQN would not be able to solve the more complex double T-maze, even though it could solve the simpler T-maze, whereas both the EC and DQN agents could solve both tasks. Note that if we decreased the number of trials to one successful trial, then the only paradigm that could potentially learn any task is EC (one-shot learning) since the other two paradigms learn incrementally.

We hypothesize that there are two main reasons why online learning is comparatively slower. First, since the agent is only rewarded where it finds the goal (sparse rewards) and updates of the DNN are based only on the current experience that connects two consecutive states, it takes many repeated experiences to propagate reward information from the goal to other states in the environment, particularly to the starting point. Second, learning is based on gradient descent on the loss function, so performing updates with single data points results in noisy updates of the DNN. This makes learning particularly unstable and accounts for the much larger variance in the performance of the online DQN agent, which is clearly visible in the learning curves for each individual run (Fig. S1). This is why mini-batch updating is prevalent in the field of machine learning to reduce the noise in gradient descent (Masters and Luschi, 2018). Hence, replaying past experiences of the agent from EM speeds up the propagation of the reward information and stabilizes learning.

The learning curves of the three agents are the main results. In the following, we analyze in more detail how and what the three agents learn to gain more insight into how EM affects learning. In particular, we test the two new hypotheses about the mechanisms by which EM speeds up learning.

### Replay learning finds better asymptotic solutions as compared to one-shot learning

To demonstrate the differences between the solutions found by the EC and DQN agents, we visualized single example trajectories that the agents took inside the tunnel maze during the test trial after 500 trials of training. A representative EC agent (Fig. 6, green arrows) took a longer trajectory than a representative DQN agent (orange arrows). Over all, 52% (13/25) of the EC agents took the trajectory along the tunnel, whereas only 12% (3/25) of the DQN agents employed the same strategy to reach the goal, i.e., the vast majority of the DQN agents found the shorter orange trajectory. We explain the difference between the agents behavior as follows. The green trajectory can be found more easily by random exploration, because once the agent moves into the tunnel, it is very likely that the agent moves along the tunnel and reaches the goal since its movement is constrained by the walls. By contrast, the shorter trajectory is harder to discover since it requires exploration of the open space at the bottom half of the tunnel maze, which takes more trials and errors. A DQN is more likely to find the better solution, because it learns more slowly and, thus, explores the environment more extensively (more on this below). The EC agent, by contrast, learns faster, but it is more prone to get stuck in a sub-optimal solution when there are multiple solutions and the globally optimal one is more difficult to find.

**Figure 6:**
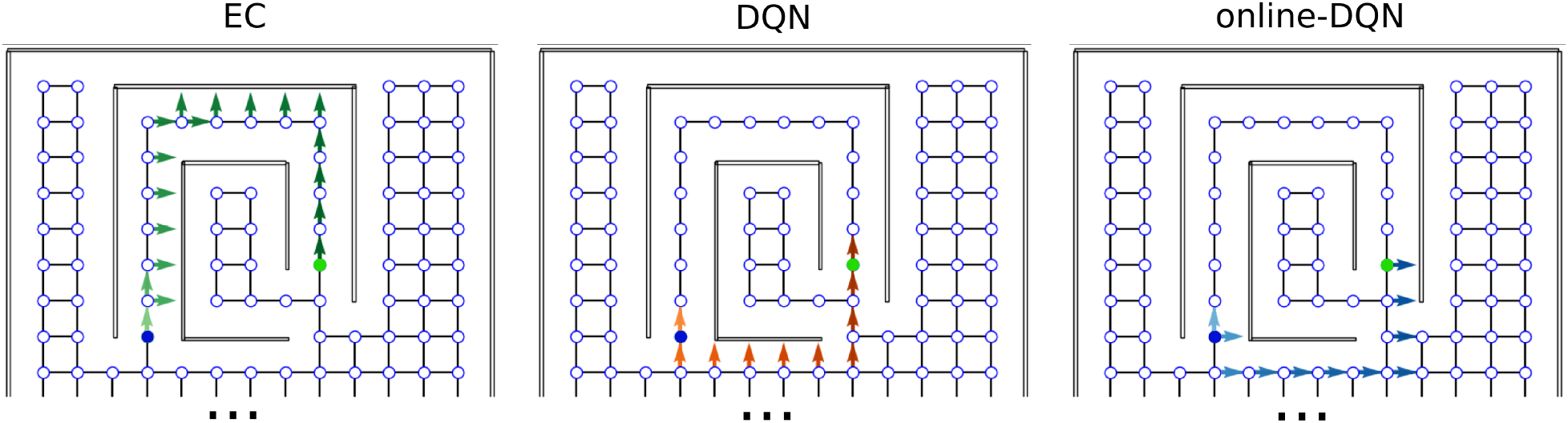
Sample trajectories in the tunnel maze for three learning algorithms. An arrow is attached to each node that the agent visits in one test trial. The direction of the arrow indicates the orientation of the agent, not the chosen action; lighter colors represent visits earlier in the trajectory and darker colors later in the trajectory. The starting and goal node are depicted in blue and green, respectively. Model-free Episodic control (EC), Deep Q network (DQN), and online Deep Q network (online-DQN).

To compare the asymptotic solutions found by the three learning algorithms more systematically, we placed the trained agents on the starting position inside each maze and recorded the number of time steps they took to find the goal (Fig. 7). This analysis revealed two interesting results. First, there were always a few simulations where the online DQN agent could not find a solution from the starting node to the goal, indicated by the data points around 600 time steps (the time-out value). It is these unsuccessful test trials that contribute to the high average number of time steps in the solutions found by the online-DQN agent. Nevertheless, even the online DQN agent discovered trajectories with lengths similar to those found by the EC and DQN agent, as indicated by the data points inside the blue bars. This confirms that the online learning paradigm is an unstable process whose success depends on the randomness in the training.

**Figure 7:**
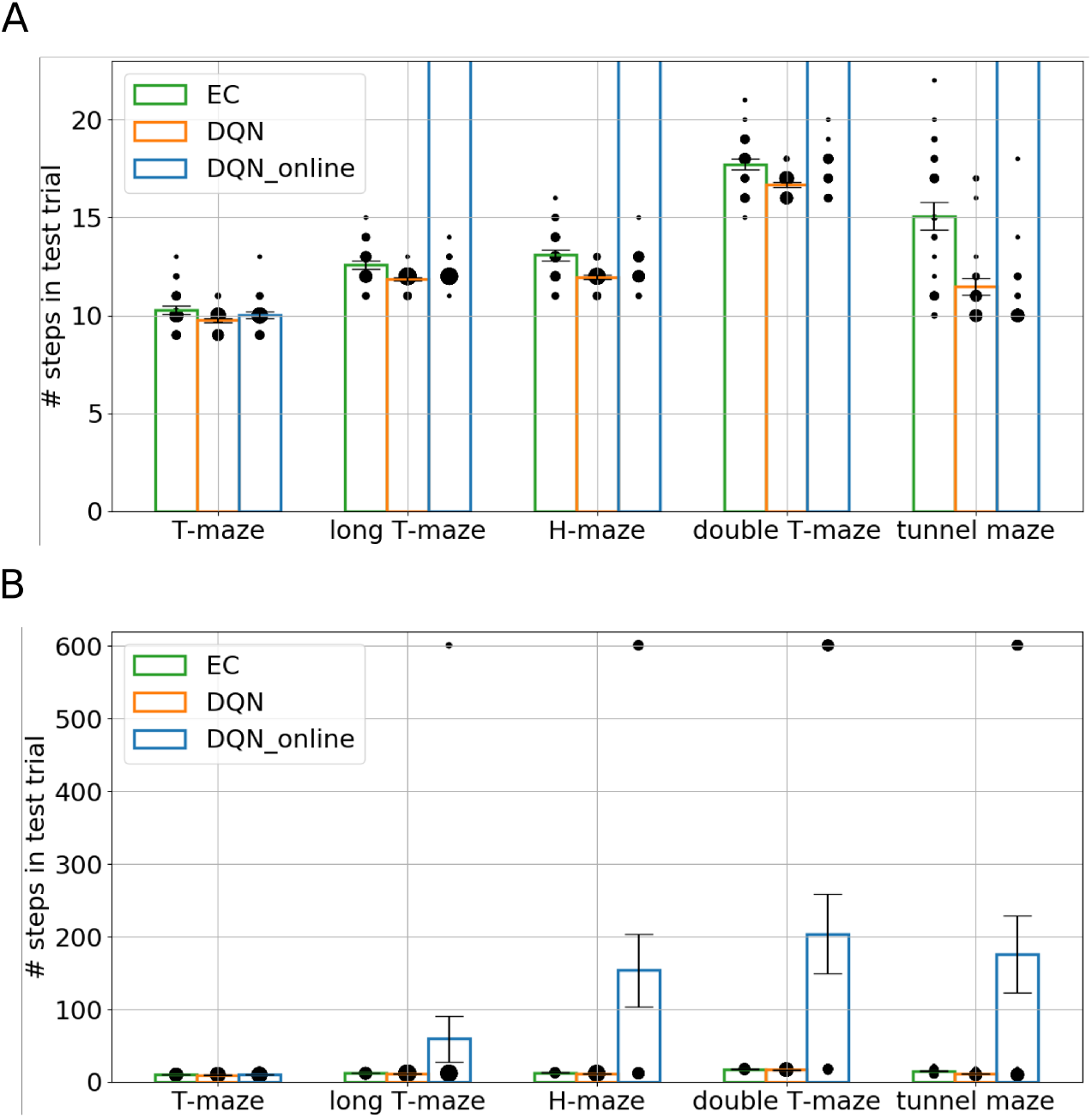
The number of time steps that the agent takes to find the goal in test trials. Bars represent the average number of steps for each algorithm-task combination over 25 runs. The size of each black dot represents the frequency of the appearance of a data value within the 25 runs. The vertical line on each bar represents the standard error of the mean. **A** and **B** show the same results with different Y-axis ranges.

Second, in every environment the DQN agent found better solutions, i.e. shorter average trajectories, than the EC agent did (Fig. 7). This difference is especially pronounced in the tunnel maze where there are multiple different routes from the start to the goal (see Fig. 4E), because of how EC and DQN utilize episodic memory. EC directly retrieves a sequence of past experiences for making decisions, and therefore follows the first solution it finds, which is discovered through randomly exploring the maze and, hence, is not necessarily optimal. By contrast, DQN constantly extracts the optimal solution based on all its past experiences, which is a gradual process, enables the agent to explore the environment more extensively, and yields more diverse experiences compared to EC. Thus, it is more likely for the DQN agent to extract a near-optimal solution. Of course, it is possible that the first solution found by the EC agent is near optimal, but the probability of this incident is small, especially when the state space is large. This also means that the solutions found by EC between independent runs can be very different, which explains the larger variance of time-steps required to reach the goal for the EC as compared to the DQN agent (green and orange bars, respectively, Fig. 7).

### One-shot learning propagates reward information more efficiently than replay learning does

Next, we examined how extensively the three agents explored the environments during training and how well the reward information was propagated from the goal to other states (Fig. 8). In the training phase, we recorded the position (represented by a node index) and orientation of the agent inside the mazes at each time step and extracted only the unique combinations to represent the set of states that the agent visited during training. Note that each state (location, orientation) is uniquely associated with an RGB image. After the training was complete, we placed the agent in every state it had visited during training and checked whether the agent would navigate to the goal from that initial state, or not. If so, we considered the reward to have propagated to that initial state and call that state a solution state.

**Figure 8:**
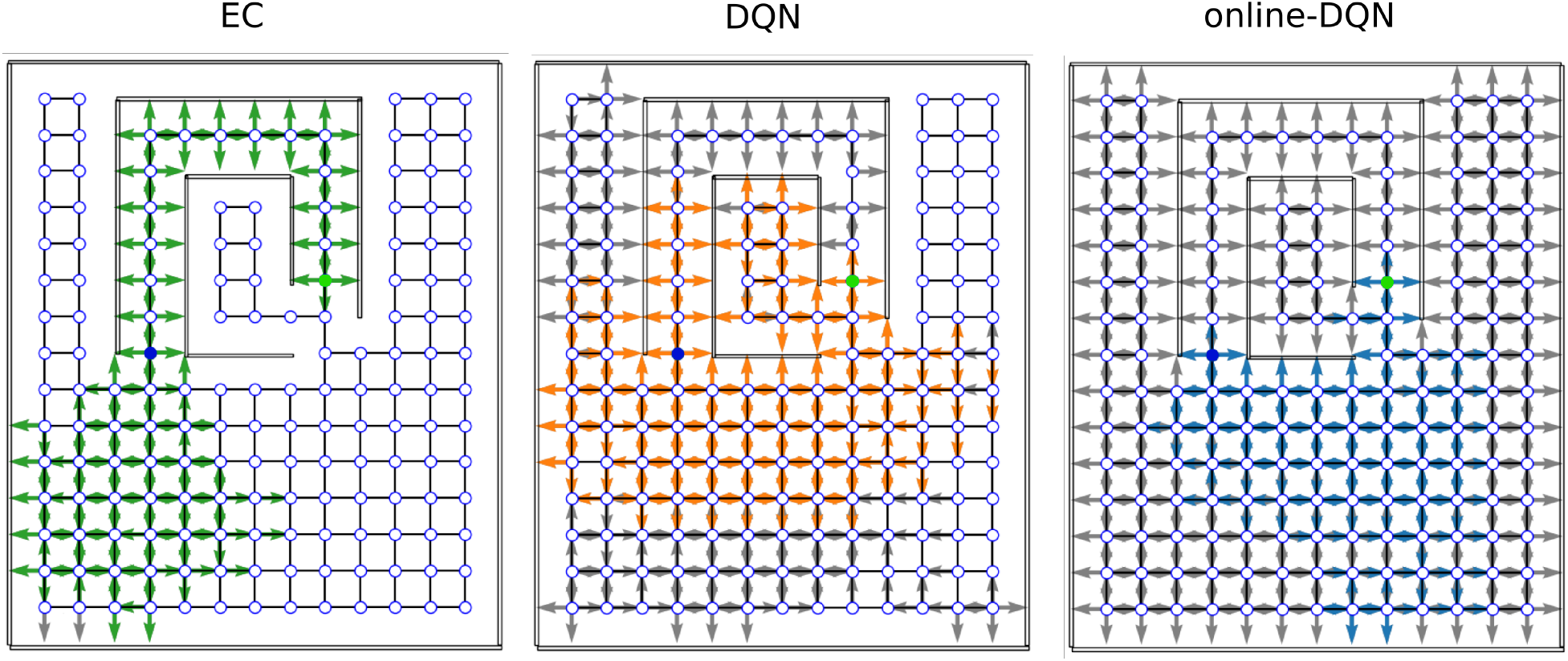
Sample coverage and propagation of reward information in the tunnel maze. An arrow (gray/colored) indicates that the agent has visited the node. The direction of the arrow indicates the orientation of the agent, not the chosen action. A colored (green/orange/blue) arrow indicates that the agent can navigate to the goal, if placed on that node in that orientation. The starting and goal nodes are depicted in blue and green, respectively. (EC: Model-free Episodic control, DQN: Deep Q network, online-DQN: online deep Q network).

In one example in the tunnel maze (Fig. 8), the EC agent explored the least, but spread the reward information to most of the states it had visited. It is also apparent that the agent always went into the tunnel to reach the goal no matter which state it starts from, since the reward information is not propagated to the node below the goal node (marked in green). The presence of gray arrows indicates that there are states that were visited by the EC agent, from where it however cannot find the goal. Since EC propagates the reward information in one shot at the end of each trial (Eq. 4), these states were probably visited in the very early stage of training when the agent failed to find the goal before the trial timed out. Finally, due to rapid learning, the behavior of the EC agent soon becomes highly repetitive after the first solution is discovered. By contrast, the DQN agent explored the maze more extensively, and also propagated the reward information to a large portion of the visited states as well. It is not easy to tell at which stage of the training the agent visits a gray state since the DQN propagates the reward signal gradually. Lastly, the online DQN agent explores the entire maze, but spreads the reward information to the smallest range out of the three algorithms. This demonstrates the advantages of memory replay as the DQN agent is able to find the goal starting from more states by exploring a smaller range of the environment compared to the online DQN.

A more systematic analysis (Fig. 9) confirms that, on average, the fractions of visited states in each maze for the three learning algorithms are inversely related to the learning speed, namely, online DQN *>* DQN *>* EC. That is, the faster the agent learns, the less it explores the environments. Although the online DQN agent explored almost 100% percent of the state space in each maze, it propagated the reward information the least among the three algorithms. There does not appear to be a significant difference in the fraction of states from where the EC and DQN agent could navigate to the goal. However, since the EC agent explores a smaller fraction of the environment, it seems to have propagated the reward information more efficiently to the states it had visited. To analyze this advantage of EC, we plotted the ratio of the number of solution states to the total number of visited states for each algorithm-maze combination (Fig. 10). Indeed, in every maze, the EC agent propagated the reward information to a larger fraction of the explored states than the DQN agent did.

**Figure 9:**
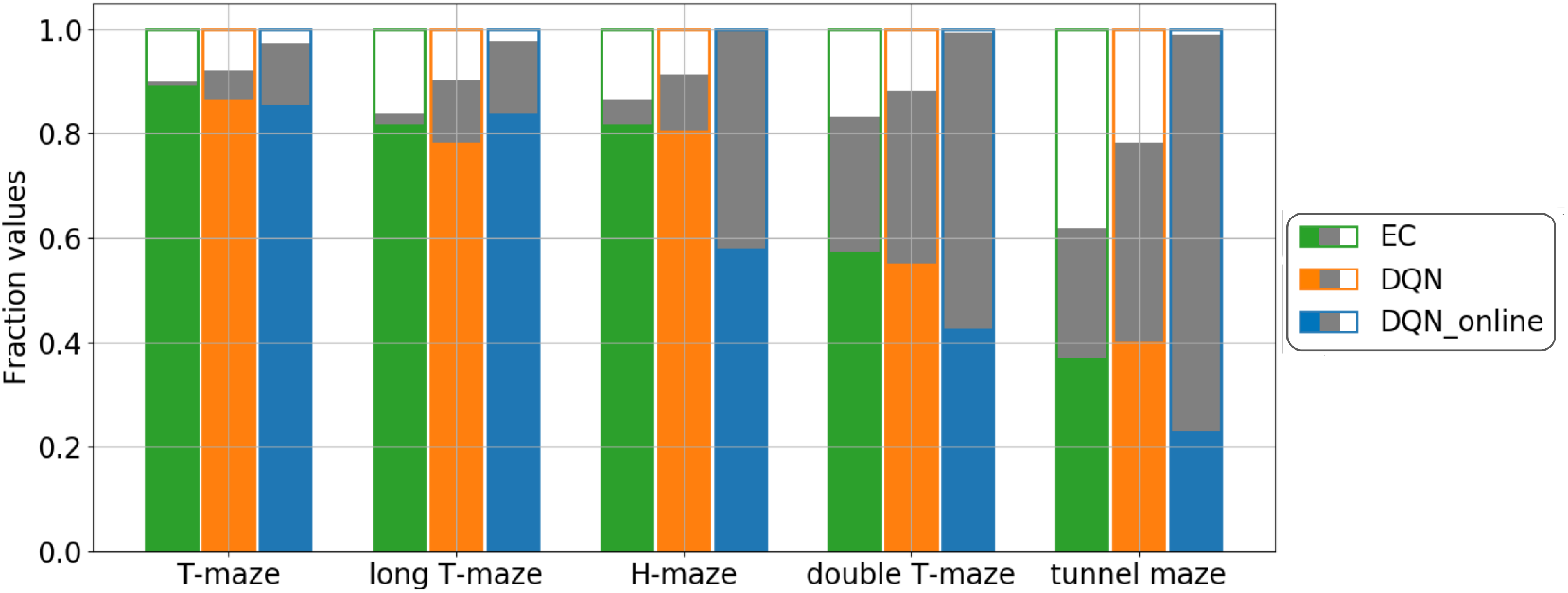
Summary of training outcomes. For each algorithm-task combination, bars show the fraction of states that were never visited during training (white fill), from which the agent will navigate to the goal, i.e., solution states (green/orange/blue fill), and were visited at least once during training, but from which the agent will not reach the goal (gray fill). The visited states were collected from the training of 500 trials and the solution states from 1 test trial after the training. The results represent an average over 25 runs.

**Figure 10:**
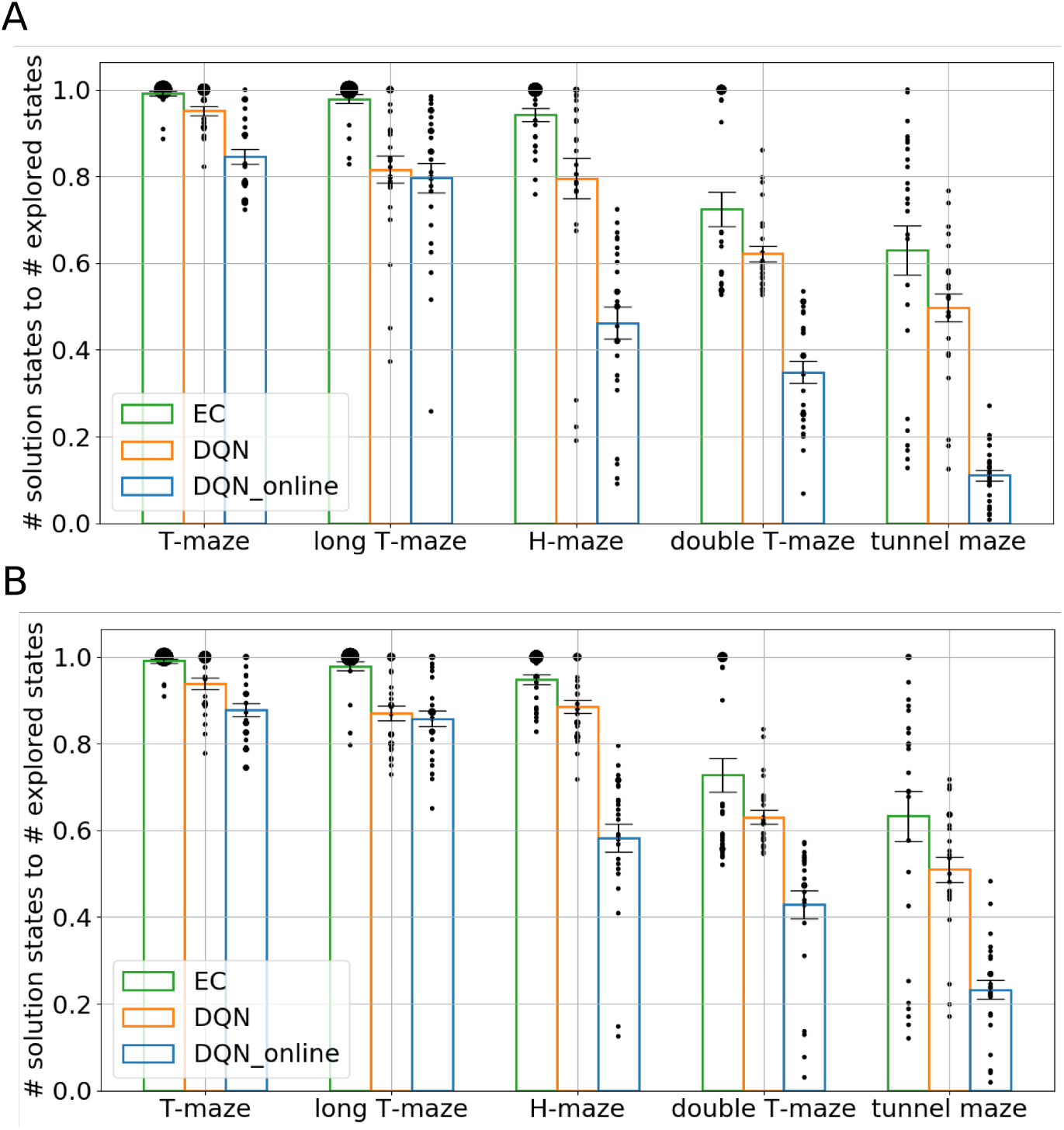
Efficiency of reward propagation. The ratio between the number of the solution states, i.e., states from which the agent will navigate to the goal, and the number of explored states during the training. Each bar represents an average over 25 simulations. The size of each black dot represents the frequency of the appearance of a data value within the 25 runs. The vertical line on each bar represents the standard error of the mean. **A**: After 250 learning trials. **B**: After 500 learning trials. While more learning trials allows replay (DQN) and online learning to propagate the reward information to a slightly larger fraction of visited nodes, the improvement is small.

Finally, we analyzed whether the DQN and online DQN agents would propagate the reward information further, if given more training trials. By comparing the training outcome after 250 trials (Fig. 10A) to that after 500 trials (Fig. 10B), we see that the solution-states-to-visited-states ratio for the EC and DQN agents have converged, but it is quite possible that the online DQN agent would spread the reward information to a larger fraction of states, if it were given more training.

### Near-optimal hybrid learning and the effect of disrupting replay while episodic memory remains intact

Animal learning in natural settings is most likely a combination of the three learning paradigms studied here. After analyzing the characteristics of each learning paradigm separately, we therefore combined the three agents in one hybrid agent (see section *Hybrid learning*). We chose the tunnel maze for the experiments and compared three agents: hybrid, EC, and DQN. The hybrid agent is able to take advantage of the strengths of both the EC and DQN (Fig. 11A), i.e., it initially learns faster than the DQN and approaches the learning speed of EC. Nevertheless, the hybrid agent learns a little more slowly than EC, because there are larger fluctuations in the output of the neural network during early training, which interferes with the values from the *Q* table that is updated by EC. The hybrid agent reaches an asymptotic performance that is as good as that achieved by the DQN agent (Fig. 11B), while learning faster than the latter. Hence, the hybrid agent combines the fast learning of EC with the better asymptotic performance of the DQN for best overall performance.

**Figure 11:**
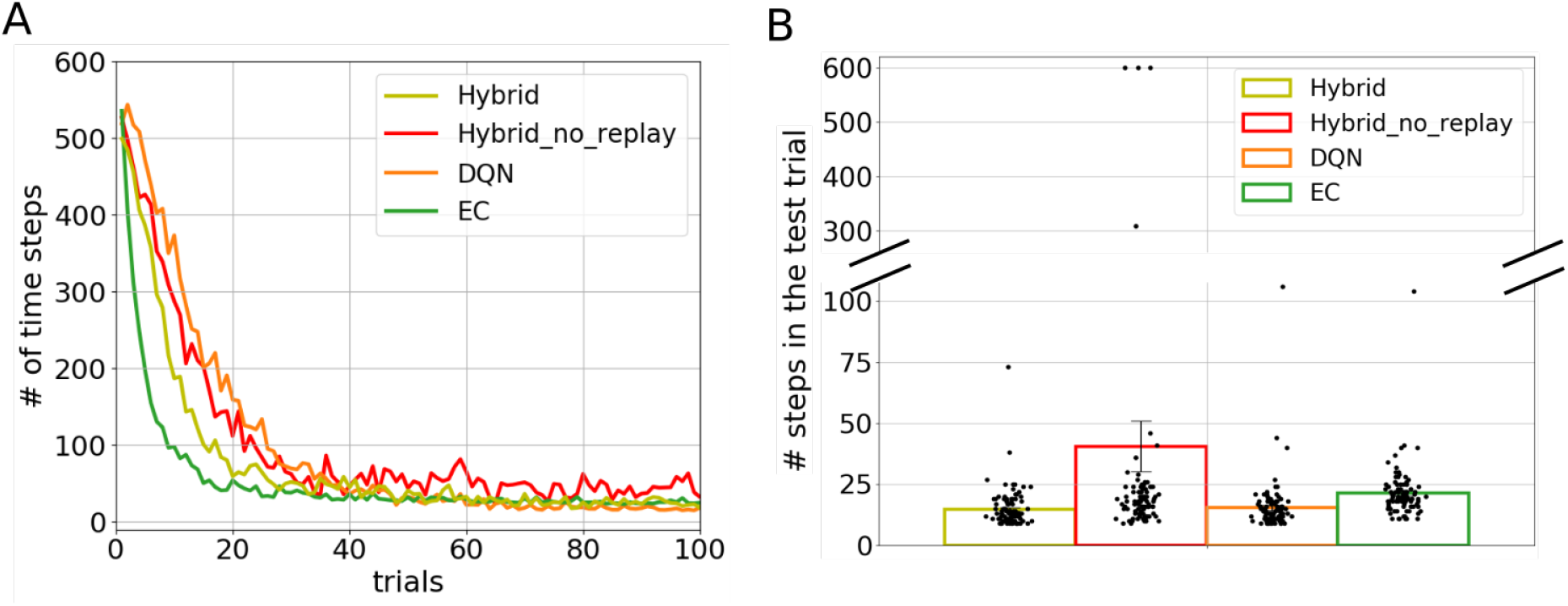
Hybrid learning combines rapid initial learning and near-optimal asymptotic performance. **A**: Learning curves, averaged over 100 independent runs, show that the hybrid agent that combines all three learning paradigms initially learns faster than the DQN, almost as fast as the EC. Even with replay impaired the hybrid agent learns as rapidly as the DQN, which uses replay, due to integration of EC (one-shot learning). **B**: Asymptotic performance of the hybrid agent is as good as that of the DQN, but suffers severely if replay is impaired. Shown are the number of time-steps that agents take to find the goal in the test trial. Bars represent the average number of steps over 100 runs. Each black dot represents a single run.

The disruption of hippocampal replay, by interfering with Sharp-wave-ripples (SWR) in the hippocampus, during the resting session of a spatial navigation task slowed down learning compared to controls (Girardeau et al., 2009; Ego-Stengel and Wilson, 2010; Jadhav et al., 2012), (Fig. 12, left). To model the effects of replay disruption, we studied a hybrid agent in which experience replay is removed from the update of the neural network. This hybrid agent without replay learned more slowly than the intact hybrid agent in the first 30 trials of training (Fig. 11A and Fig. 12, right), consistent with experimental observations. Note that if when we limited the number of simulations to be the same as the number of rats used in the experiment (*N* = 6) there was substantial variance in the finding (Fig. S2). However, the trend is consistent. The hybrid agent without replay also reached a worse asymptotic performance and had a much higher variance compared to the other three agents (Fig. 13). This result is in accordance with the higher variance of the online DQN agent, discussed above. Our modeling result is also consistent with experimental observations that replay disruption over an extended period prevented animals from fully learning a spatial task (Girardeau et al., 2009).

**Figure 12:**
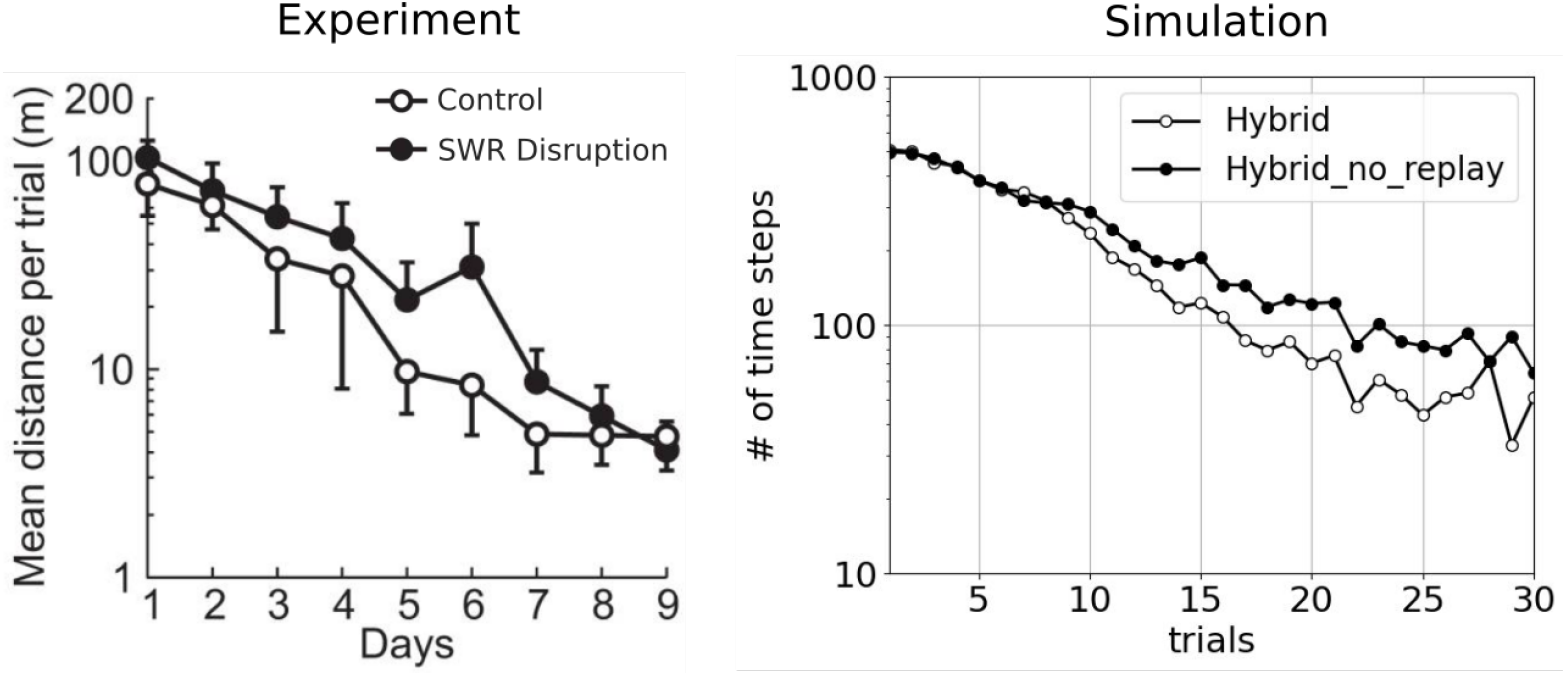
Simulations reproduce experimental results of replay suppression. **Left**: Results adapted from Ego-Stengel and Wilson (2010) where the animals need to navigate from a fixed starting point to a fixed goal inside a maze. Filled dots represent the group of rats where sharp-ware-ripples were disrupted during post-task rest. *N* = 6. **Right**: Results from our simulation of the hybrid agent when replay was impaired qualitatively match the experimental results. Averaged over 100 simulations.

**Figure 13:**
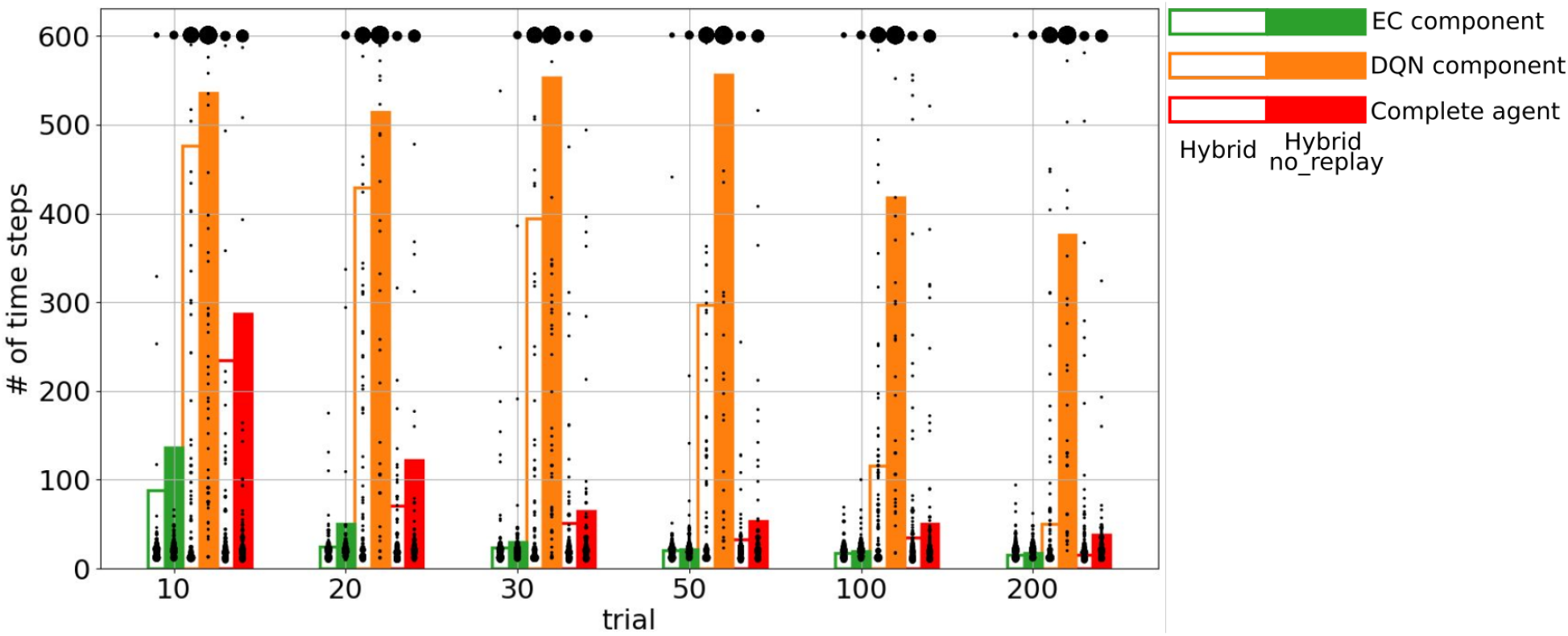
Performance of the EC component, DQN component and the complete hybrid agent at different training stages. Empty bars represent the results for the hybrid agent with memory replay, filled bars the hybrid agent without replay. Bars indicate the average number of steps over 100 runs. Each black dot represents a single run and the size of the dot represents the frequency of the data point. The larger the dot is, the more frequently the corresponding value appears among the 100 runs.

To see why removing memory replay impedes learning speed and asymptotic performance of the hybrid agents, we examined the learning profiles of the EC component (*Q* table) and the DQN component (*Q* network) separately at different stages of learning (Fig. 13). To this end, we disabled one of the components in the model and examined the performance of the other. The most striking difference exists between the solutions found by the DQN components with and without replay, especially at late stage of the training (> 50 trials). This result makes sense since memory replay speeds up and stabilizes the learning of the Q network dramatically, as discussed in previous sections, and the difference between replay and no replay becomes larger the more experiences there are that can be replayed. By contrast, the EC components in the hybrid agents show the opposite pattern. The largest difference between replay and no replay are observed during the initial stages of training and it attenuates quickly within < 30 trials. Nevertheless, the EC components in both hybrid agents learn in a one-shot manner and their performance is already quite high early on.

The initial difference between the two EC components must have arisen from differences in how the environment was explored, which is consistent with our finding that the online DQN agent explores the environment more broadly (Fig. 8).

Taken together these observations account for the initial learning speeds and asymptotic performance of the two hybrid agents. Since the hybrid agents choose the larger Q value from the EC or DQN components to select the next action, good performance of one component will drive good overall performance of the hybrid agent, hence ignoring the weak performing component. So, both hybrid agents initially learn quickly because the EC component does regardless of the DQNs’ weak performance, and the difference between the hybrid agent with and without replay can be attributed to the difference between their EC components. Also, the asymptotic performance of the hybrid agent with replay is driven by the good asymptotic performance of its DQN component. However, if the weak component has large variability in its outputs, it can act as a source of noise which interferes in the decision-making and, hence, impair the overall performance of the hybrid agent. So, the hybrid agents initially learn slightly slower than the pure EC agent because of interference from the DQN. Inference also impairs the asymptotic performance of the hybrid agent without replay, because its DQN component fails to converge.

## Discussion

To investigate the functional role of episodic memory (EM) in spatial learning, we studied three learning paradigms that differ in how they access information stored in EM: one-shot learning, replay learning and online learning. To compare the three paradigms quantitatively, we chose three reinforcement learning algorithms and applied them to spatial learning tasks in simulated maze environments. The three agents received no prior information about the mazes and had to solve the navigation tasks based on raw sensory inputs by trial and error. We found that whether an agent is able to learn the task depends on the number of learning trials and the complexity of the task. One-shot learning initially solves the task very quickly, but cannot reach the same asymptotic performance as replay learning, which converges more slowly but explores the environments more extensively. Online learning without EM is the most sensitive to changes in task complexity and is unstable. It therefore leads to large variability in learning performance. Furthermore, we showed that it is possible to combine all learning paradigms in a hybrid agent such that it benefits from both fast learning and best asymptotic performance. Removing replay from this hybrid agent shows the same effects on learning and asymptotic performance as animals in which hippocampal replay was selectively disrupted. We further discuss the implications of our results below.

### Implications for the role of the hippocampus in learning

Many tasks in animal experiments have been categorized as being hippocampally dependent, such as, e.g., spatial navigation (Morris et al., 1982), contextual fear conditioning (Maren et al., 2013), and trace conditioning (McEchron et al., 1998), because hippocampal animals perform worse in these tasks than controls. However, the literature suggests that the hippocampus might play diverse roles in these different tasks. In spatial navigation, the most widely held view suggests that the hippocampus represents a cognitive map (Moser et al., 2008), which allows animals to localize themselves and plan trajectories to a goal location. However, the same brain region is also thought to be responsible for constructing a contextual representation in contextual fear conditioning (Maren et al., 2013), or bridging a temporal gap between the CS and US in trace conditioning (Bangasser et al., 2006).

Our current modeling results suggest a rather different picture. First, it might not be useful to categorically label a task as hippocampally dependent, or not, since the dependence is determined by the complexity of the task and how extensive the training is. For instance, a simple spatial navigation task might not depend on the hippocampus, if a sufficient number of training trials are given to the hippocampal animal, which can still use online learning. Second, a hippocampal lesion might have more widespread impact on behavior than traditional measures of success suggest. In our simulations, a hippocampal lesion not only significantly decreases the learning speed, but also leads to a more thorough exploration of the environment, as well as, more variability in the learning process and outcomes due to the high instability of online learning. Third, the function of the hippocampus is to serve as a crucial part of the biological substrate of the EM system (Cheng, 2013; Cheng and Werning, 2016), so that animals can perform one-shot learning and replay learning by accessing EM in two different modes. We suggest that all other cognitive functions that the hippocampus might be involved in can be traced back to its function in EM.

### Two different modes of accessing episodic memory

To the best of our knowledge, ours is the first study that modeled and quantitatively compared two different modes of accessing EM. On the one hand, retrieval entails the direct use of episodic memory and supports one-shot learning. It has been observed and discussed in both animal and human experiments (Öhman et al., 1975; Steele and Morris, 1999; Tse et al., 2007). Recently, Banino et al. (2020) have shown that retrieving EM with a recurrent attention mechanism enables transitive inference. On the other hand, hippocampal replay has been thought to play a role in memory consolidation for decades (Buzsaki, 1989; McClelland et al., 1995). It has been demonstrated in computational models (McClelland et al., 1995; van de Ven et al., 2020) that replay can prevent catastrophic interference (McCloskey and Cohen, 1989; Kirkpatrick et al., 2017) in semantic networks by enabling interleaved training, i.e., the alternating presentation of old and new information. This latter aspect of replay might be more important than the mere repetition of the experience and explains why increasing the learning rate in the online learning paradigms does not improve performance to the level of replay learning. However, the difference between these two operating modes of EM has not been studied.

In addition, we view the function of memory replay differently from the standard model, which suggests that hippocampal replay transfers episodic contents from a hippocampus-dependent format to one that involves neocortex alone. Alternatively, we suggest that memory replay enables the neocortex to extract statistical regularities from the stored experiences, and that the information represented by the neocortex in the end is different from the information in EM Cheng (2017).

In our simulations, the two modes of accessing EM learn at different speeds and, as a consequence, lead the agent to explore the environment to different extents. Particularly, replay learning enables the agent to find better asymptotic solutions since the agent explores a larger fraction of the state space. By contrast, the one-shot learning agent shows highly repetitive behavioral patterns after finding the solution for the first time. Therefore, our results suggest that impairing replay, but leaving EM intact otherwise, will lead to very specific learning deficits that are different from deficits due to abolishing EM altogether, i.e., anterograde amnesia. However, it is not easy to observe this distinction at the behavioral level in animal experiments since the two modes coexist in a healthy brain and are normally both impaired by a hippocampal lesion.

We therefore studied a hybrid agent in which all three learning paradigms were combined as a model of a control animal. This enabled us to dissociate the functional role of replay from that of EM in general by removing replay from the hybrid agent while leaving EM, and hence one-shot learning, intact. These studies qualitatively reproduced the slower learning and lower asymptotic performance in animals when SWRs in the hippocampus were disrupted to prevent the replay of place cell activity (Girardeau et al., 2009; Ego-Stengel and Wilson, 2010). Further analysis revealed that the behaviour of the hybrid agent is primarily driven by the component with the better performance, as intended. However, if the weaker component exhibits large variability in its output, it can act as a source of noise, interfering with the overall performance of the hybrid agent. Our modeling results suggest that disrupting replay affects learning speed and performance because it impairs memory consolidation, which increases the variance in the output from the semantic system, and in turn interferes with the direct use of episodic memory when the animal makes a decision. We therefore predict that one-shot learning could be recovered in replay disruption experiments, if the influence of the semantic network could be suppressed during decision making.

### On the importance of studying the learning dynamics

Our modeling in this study suggests that it is critical to experimentally measure the learning curves in simple and more complex tasks in control and hippocampal animals, as well as, when hippocampal replay or memory retrieval is inhibited separately. While in hippocampal research it is fairly common to report learning curves, they are generally averaged over animals and/or blocks of learning. However, such averaging can produce misleading results (Gallistel et al., 2004; Smith et al., 2004). For instance, Cheng and Sabes (2006, 2007) show that studying the dynamics of learning reveals an enhanced picture of the process of adaptation, and Donoso et al. (2021) uncovered a large variability of behavior in extinction learning and renewal by focusing on individual learning curves. None of these findings could have been revealed by simply comparing the differences in performance before and after learning, or if data was averaged across individuals. Studying individual learning curves reveals much more about the learning process and the role of episodic memory, than comparing pairs of data points, e.g., comparing the performance of hippocampal to control animals in one task after a fixed number of learning trials. Hence, our modeling provides another example in which much more information can be gained from studying the trial-by-trial dynamics of learning in individuals.

### The sequentiality of episodic memory

While there are exceptions (Levy, 1996; Cheng, 2013; Bayati et al., 2018), sequentiality has not been widely accepted as an important feature of EM and many models have represented EM as static memory patterns (Treves and Rolls, 1992; Hasselmo et al., 2002; Káli and Dayan, 2004). In our model, the sequentiality of episodic memory is key in the one-shot learning paradigm. This is because when there is a temporal distance between the agent’s actions and their consequences, the sequential order of past experiences has to be maintained in memory to credit the past states and actions when updating the Q values. The importance of sequentiality is consistent with recent observations that theta sequences in the hippocampus are modulated by the animal’s previous behaviour at the spatial location where the theta sequences occur (Parra-Barrero et al., 2021).

Although an agent is still able to extract semantic knowledge through memory replay without maintaining the sequential order, its learning speed is relatively slower and it cannot solve a task, especially a difficult one, in a one-shot manner. However, since life-threatening experiences should not be repeated, one-shot learning is crucial for the survival of an animal or species. In turn, one-shot learning in a complex task requires that sequences of experiences are stored in EM.

### The role of sequential replay

In our model, experiences are replayed individually in random order. While in the hippocampus, although activity patterns might sometimes be replayed in random order. they are also replayed in sequential order (Louie and Wilson, 2001; Buhry et al., 2011). We did not model a biologically-plausible replay mechanism because, first, an artificial neural network is notoriously hard to train with correlated data (Bengio, 2012), and replaying training samples randomly breaks the serial correlations among the data, thus stabilizing the learning. Therefore, in the original implementation of deep Q learning (Mnih et al., 2015), experiences are randomly replayed and we chose to leave this part unchanged in our modeling. Second, the statistics of hippocampal replay in rodents is complex (Gupta et al., 2010; Ólafsdóttir et al., 2015; Stella et al., 2019) and difficult to model. Since the primary focus of this paper is about the functional consequence of memory replay, not the statistics of replay *per se*, we decided to simplify our modeling at the current stage. However, we believe there is a functional role of the sequential replay in the hippocampus and will investigate it in future work.

In conclusion, our modeling study made a first attempt at studying the computational role of EM in spatial learning and quantitatively comparing two different modes of accessing EM: retrieval and replay. While many open questions remain, the computational framework allows us to clearly delineate episodic from semantic information and therefore could help resolve the controversial issue of what, if anything, separates episodic from semantic memory.

## Materials and Methods

### Computational modeling framework

All simulations were performed in a virtual-reality modeling framework (Fig. 2) that was developed to study models of rodent behavior in spatial navigation and extinction learning (Walther et al., 2021). This framework is named CoBeL-RL (**C**losed-l**o**op simulator of complex **Be**havior and **L**earning based on **R**einforcement **L**earning, https://doi.org/10.5281/zenodo.5741291). The virtual environments were designed with the unity game engine (https://unity.com/), while the remaining parts of the framework were developed using Python. In the simulations, an artificial agent equipped with a wide-view camera represented an animal navigating in the virtual environment. The World Topology module determined the spatial positions and orientations, which the agent could be placed in, and the allowed transitions between them. The OpenAI Gym Interface (https://gym.openai.com/) module formed the interface between the virtual environments and the RL agent, transmitting information and control commands between the two.

### Simulated spatial learning tasks

We designed a series of virtual environments for spatial navigation in the Unity simulator to test our hypotheses (Fig. 4A-E). The agent was always placed in the same starting location (blue square) and had to find a goal (green disk). The sensory input to the learning agent consisted of an 84 × 84 RGB image (e.g. Fig. 4F). The topology graph of each environment is only used to determine which transitions are valid in the simulation and are not known to the learning agent. The agent could face four different directions at each node: north, west, south, or east; it could take six different actions: go forward, go backwards, go left, go right, rotate 90 degrees clockwise, or rotate 90 degrees counterclockwise. The complexity of the spatial navigation tasks increases from Fig. 4A to Fig. 4E, because both the total number of nodes and the minimum number of transitions that the agent needs to reach the goal increase. Once the agent reached the goal location, or the trials timed out after 600 time steps, we ended the current trial and returned the agent to the initial position. At the beginning of the experiment, the agent had no prior knowledge about the environment, hence it could only explore randomly at the beginning of training. To reduce the variance caused by the randomness in the training, we averaged performance measures over 25 independent runs for each task-algorithm combination where the algorithms is either EC, DQN, or online-DQN, defined below. For experiments with hybrid agents (defined below), we performed 100 independent runs since the effect of abolishing replay was more subtle and required more simulations to reveal a consistent pattern.

### Reinforcement learning (RL)

We modeled spatial learning in the framework of RL, where an agent interacts with an environment. At each time step *t*, the agent observes the state of the environment represented by *s*_*t*_ ∈ *S*, where *S* is the set of all possible states, and, in response, takes an action *a*_*t*_ *A* ∈ (*s*_*t*_), where *A*(*s*_*t*_) represents the set of all possible actions the agent can take in state *s*_*t*_. These actions, in combination with the dynamics of the environment, lead to a new state in the environment at the next time step *s*_*t*+1_, which the agent also observes. In addition, the agent receives a reward *r*_*t*+1_. These steps are repeated until a terminal state is reached, which indicates that episode (or a “trial” in the convention of neuroscience) has ended. For the spatial navigation tasks (Fig. 4), the state was represented by the RGB image that the agent received from the camera. The agent received a reward *r*_*T*_ = +1.0, if it found the goal, which ended the episode. *T* indicates the last time step of the current episode. No reward was given for any other action (*r*_*t*_ = 0 for all *t* < *T*).

In RL, the behavior of the agent and learning is driven by the discounted cumulative reward:

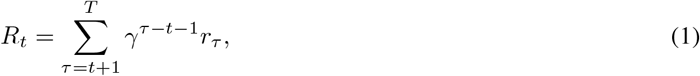

where *γ* (0 ≤ *γ* ≤ 1) is a discount factor that determines the relative values of immediate reward vs. those that are more distant in time. The objective of the agent is to learn a policy *π* that maximizes the expected cumulative reward

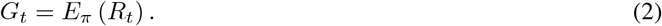

One method for solving this task is *Q* learning, a class of RL algorithms, where the agent learns a state-action value function, the so-called *Q* function. The scalar function *Q*(*s*_*t*_, *a*_*t*_) measures how desirable the state-action pair (*s*_*t*_, *a*_*t*_) is to the agent. If learned correctly, the larger the value of the *Q* function, the larger a cumulative reward the action *a*_*t*_ in state *s*_*t*_ will yield. Mathematically, the *Q* function can be expressed as

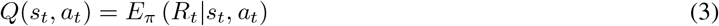

In state *s*_*t*_, the agent selects the action *a*_*t*_ with the highest *Q* value. To balance exploration and exploitation, we used the *ϵ*-greedy algorithm: with probability *ϵ* the agent will randomly select an action from *A* regardless of the *Q* values, otherwise the agent selects the action that yields the highest *Q* value. Throughout our simulations we set *ϵ* = 0.1. The discount factor is set as *γ* = 0.9 to encourage the agent to find the shortest path from the initial position to the goal.

### The three learning algorithms

We selected three RL algorithms based on *Q* learning to model our hypothesized learning paradigms (Fig. 3): Model-free Episodic Control (EC) (Blundell et al., 2016) for one-shot learning, Deep Q Learning with memory replay (DQN) (Mnih et al., 2015) for replay learning, and online Deep Q Learning (online DQN) for online learning. Real experiences were modeled as a sequence of (state, action, next state, reward) tuples, i.e., (*s*_*t*_, *a*_*t*_, *s*_*t*+1_, *r*_*t*+1_). Like in EM (Cheng and Werning, 2016), these sequences of events were stored in memory for one-shot learning in EC and for replay learning in the DQN algorithm.

#### Model-Free Episodic Control (EC)

We use *Q*^*EC*^ to refer to the *Q* function learned by Model-Free Episodic Control. *Q*^*EC*^ is represented as a table whose entries are directly computed using sequences of past experiences stored in memory by using the following equation:

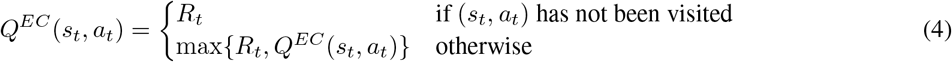

Note that the sequential ordering of experience tuples is critical in EC because *R*_*t*_ is calculated for the particular sequence in the current episode by using Eq. 1. When making a decision, the agent selects the action with the highest *Q*^*EC*^ value in the current state.

The max operation in Eq. 4 guarantees that the sequence starting from (*s*_*t*_, *a*_*t*_) that leads to the largest cumulative reward encountered so far will be followed by the agent. For states that have never been visited before, the *Q*^*EC*^ values are approximated by averaging the *Q*^*EC*^ values of the k-nearest neighbours of the given state. We used *k* = 5. Blundell et al. (2016) originally utilized two different dimension-reduction methods to pre-process the raw input in order to decrease the computational requirements: Variational Autoencoder (VAE, Kingma and Welling, 2014) and random projection. We chose the latter in our implementation. Specifically, raw images generated from Unity of size 84 × 84 × 3 were projected onto a lower-dimensional space, i.e.*ϕ, f* : *x* → *Mx* where *M* ∈ ℝ^*F* × *D*^ and *D* is the dimension of the original inputs. The entries of matrix *M* were drawn from a standard Gaussian and according to the Johnson-Lindenstrauss Lemma (Johnson and Lindenstrauss, 1984), this transformation preserves relative distances in the original space. In our implementation, the dimension of the projected space was *F* = 256. To further speed up inference and learning, the states stored in memory were used to construct a *KD* tree (short for K-dimensional tree, a binary tree where every node is a k-dimensional point) (Bentley, 1975) so that the search of closest neighbours to a given state (measured under Euclidean distance) becomes efficient. Lastly, we set the maximum number of experiences that can be stored to 50, 000.

#### Deep Q network (DQN)

The DQN represents the *Q* function as an artificial deep neural network (DNN), which maps a state *s*_*i*_ to the *Q* values of all the possible actions on this state. During learning, a mini-batch *B* of experience tuples are randomly selected from memory and used to construct a loss function according to

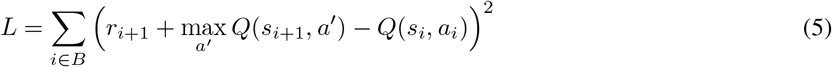

The backpropagation algorithm is used to compute the gradients of this loss function w.r.t. the weights *w* of the DNN, which are updated incrementally by applying gradient descent:

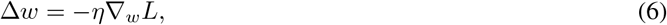

where *η* represents a learning rate. We consider the DNN to loosely represent the semantic network in the neocortex (Kriegeskorte, 2015), and the whole learning process as replaying past experiences from episodic memory to the semantic system for extracting information. Unlike for EC, the sequential ordering of experience tuples is not important for replay in DQN, because the items are randomly chosen for replay and the learning is based on the immediate reward *r*_*t*+1_ instead of the cumulative reward *R*_*t*_.

We used a DNN with the same architecture as Mnih et al. (2015). The input, an 84 × 84 × 3 RGB image, was first passed to a convolutional layer consisting of 16 8 × 8 filters with stride 4 and then to a second convolutional layer consisting of 32 4 × 4 filters with stride 2. The last hidden layer consisted of 256 fully connected units. The output layer had 6 units, each corresponding to the *Q* value of one of the possible actions. A rectifying linear unit (ReLU) was used as activation function in all layers except for the output layer, where a linear activation function was used. At each time step, a mini-batch of 32 samples was randomly drawn from the memory to update the network using the Adam optimizer with a learning rate of 0.0001.

#### Online Deep Q network

To model spatial learning without EM, online learning was based on the experiences only as they occur, i.e., each experience tuple was used only once for learning. Specifically, at each time step *t*, a loss function was constructed using only the current experience tuple according to Eq. 7. The DNN then minimized this loss function

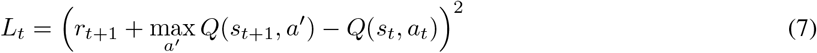

incrementally to find the optimal *Q* function. The hyper-parameters and update rules for the online DQN agent were the same as those for the DQN agent except that there was no experience replay.

### Hybrid learning

A healthy animal can both retrieve EM and replay EM as well as learn from online experiences, i.e., the three learning paradigms work in parallel and potentially interact. Therefore, we developed a hybrid learning agent in which Model-Free Episodic Control (EC) and Deep Q learning run and perform inference in parallel. The agent possessed both a table and a neural network to represent the *Q* function, where the former was updated by EC (*Q*^*EC*^) and the latter was trained with both memory replay and online experiences like the DQN and online-DQN are (*Q*^*NN*^). Specifically, the neural network was updated twice every time step, once from the current experience and the other from the replayed experiences from the memory. The agent based its decisions on the best solutions that was available, i.e.,

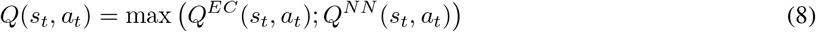

We chose the maximum operation instead of using other methods, such as weighted sum for two reasons. It does not depend on additional parameters that would require tuning and, more importantly, action selection is based on the maximum of the *Q* function. Thus, combining *Q* values in the same way is consistent with the winner-take-all flavor of the algorithms.

We decreased the number of training trials from 500 to 200, compared with previous simulations, because we wanted to compare the hybrid agent to EC and DQN and performances of these agents have converged by 200 trials.

## Acknowledgements

This work was supported by the Deutsche Forschungsgemeinschaft (DFG, German Research Foundation), project number 419037518 – FOR 2812, P2.

## Supplementary figures

**Figure S1:**
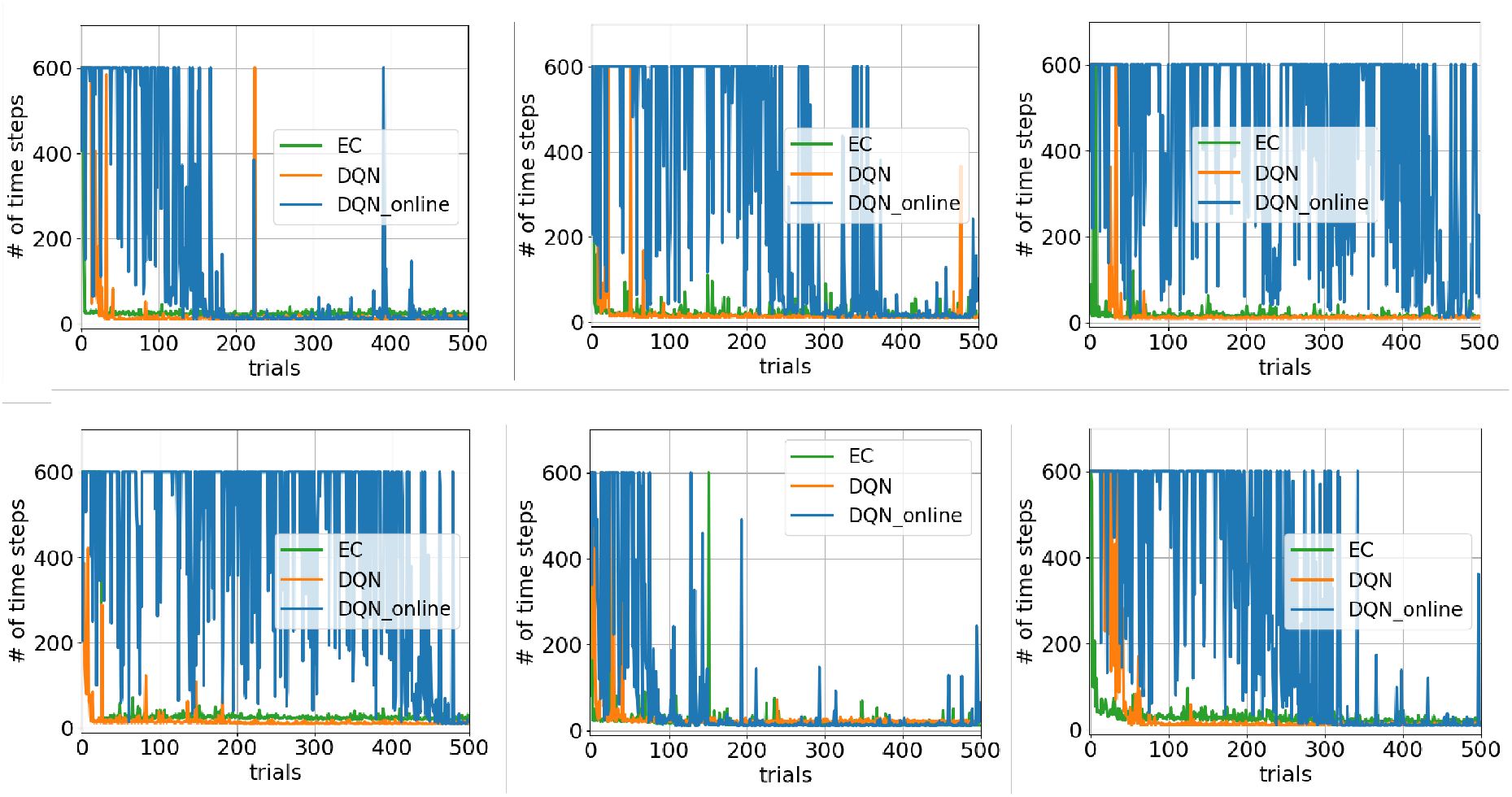
Examples of the individual learning curve for the three learning paradigms in the tunnel maze environment. Each panel shows the number of time steps that the agent takes to find the goal across trials for one agent. Six agents were selected randomly. While EC and DQN with memory replay can find the goal stably in each run, the learning for online DQN is unstable and unreliable, i.e. it does not always converge and occasionally the performance drops after convergence had been achieved.

**Figure S2:**
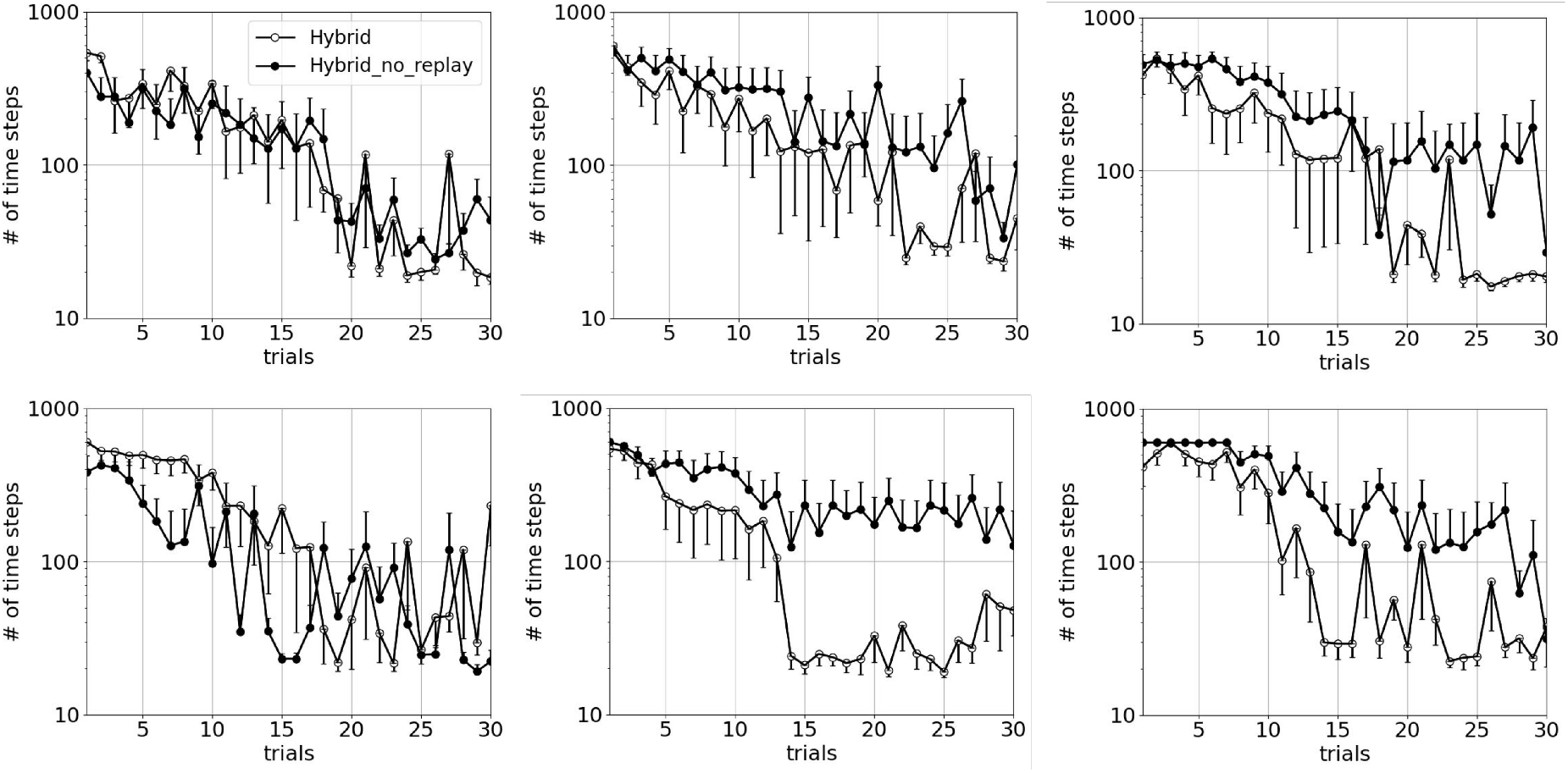
Examples of the average learning curve for the two hybrid agents in the tunnel maze for a limited number of simulations. To more directly compare our simulation results to experimental results, we performed simulations for and averaged across the same number of agents as the number of rats used in Ego-Stengel and Wilson (2010). Each panel represents the average results from *N* = 6 independent runs. The error bars represent the standard errors of the mean. While the hybrid agent with memory replay does not always learn faster than the agent without replay, there is a consistent pattern that replay benefits learning.

